# Plaat1l1 controls feeding induced NAPE biosynthesis and contributes to energy balance regulation in zebrafish

**DOI:** 10.1101/2023.12.31.573561

**Authors:** Zahra Mashhadi, Linlin Yin, Noura S. Dosoky, Wenbiao Chen, Sean S. Davies

## Abstract

Dysregulation of energy balance leading to obesity is a significant risk factor for cardiometabolic diseases such as diabetes, non-alcoholic fatty liver disease and atherosclerosis. In rodents and a number of other vertebrates, feeding has been shown to induce a rapid rise in the intestinal levels of *N*-acyl-ethanolamines (NAEs) and the chronic consumption of a high fat diet abolishes this rise. Administering NAEs to rodents consuming a high fat diet reduces their adiposity, in part by reducing food intake and enhancing fat oxidation, so that feeding-induced intestinal NAE biosynthesis appears to be critical to appropriate regulation of energy balance. However, the contribution of feeding-induced intestinal NAE biosynthesis to appropriate energy balance remains poorly understood because the specific enzymes responsible for feeding-induced NAE biosynthesis have not been identified. The rate-limiting step in the intestinal biosynthesis of NAEs is formation of their immediate precursors, the *N*-acyl-phosphatidylethanolamines (NAPEs), by phosphatidylethanolamine *N*-acyltransferases (NATs). At least six NATs are found in humans and multiple homologs of these NATs are found in most vertebrate species. In recent years, the fecundity and small size of zebrafish (*Danio rerio*), as well as their similarities in feeding behavior and energy balance regulation with mammals, have led to their use to model key features of cardiometabolic disease. We therefore searched the *Danio rerio* genome to identify all NAT homologs and found two additional NAT homologs besides the previously reported *plaat1*, *rarres3*, and *rarres3l*, and used CRISPR/cas9 to delete these two NAT homologs (*plaat1l1* and *plaat1l2*). While wild-type fish markedly increased their intestinal NAPE and NAE levels in response to a meal after fasting, this response was completely ablated in *plaat1l1^-/-^ fish.* Furthermore, *plaat1l1^-/-^*fish fed a standard flake diet had increased weight gain and glucose intolerance compared to wild-type fish. The results support a critical role for feeding-induced NAE biosynthesis in regulating energy balance and suggest that restoring this response in obese animals could potentially be used to treat obesity and cardiometabolic disease.

## INTRODUCTION

Dysregulation of energy balance leading to obesity is one of the key risk factors for cardiometabolic diseases including type 2 diabetes (T2D), non-alcoholic fatty liver diseases (NAFLD), and cardiovascular disease (CVD) (1–4). The regulation of various aspects of energy balance including feeding behavior and metabolism is highly complex and includes multiple signaling mechanisms (e.g. CCK, GLP-1, insulin, leptin). Recent data suggests that *N*-acyl-ethanolamines (NAEs) including *N*-palmitoyl-ethanolamine (PEA), *N*-stearoyl-ethanolamine (SEA), *N*-oleoyl-ethanolamine (OEA), and *N*-linoleoyl-ethanolamine (LEA) may also play a critical role in energy balance. Previous studies have demonstrated that when lean fasted rodents consume a meal, there is a rapid rise in the intestinal levels of OEA and other NAEs (5, 6). Similar feeding-induced rises in NAEs have been observed in pythons(7), and goldfish(8). Administration of NAEs including OEA, PEA, SEA, and LEA exert biological effect considered important in regulating energy balance and protecting against cardiometabolic diseases such as release of GLP-1(9), inhibition of food intake(10–13), increasing expression of fatty acid oxidation genes, and enhancing resolution of inflammation(14, 15). In rodents, chronic consumption of high fat diet (HFD) results in the loss of feeding-induced NAE biosynthesis(10, 16–18). Together, these results suggest that feeding-induced NAE biosynthesis plays a critical role in energy balance, so that the loss of feeding-induced NAE biosynthesis may be an important contributor to energy balance dysregulation and obesity. However, because the specific enzymes responsible for feeding-induced NAE biosynthesis have not been identified, the contribution of feeding-induced NAE biosynthesis to appropriate energy balance remains poorly understood.

The rate limiting step in the formation of NAEs is the formation of their precursor *N*-acyl-phosphatidylethanolamines (NAPEs) (19). These are generated by the transfer of an *O*-acyl chain from a donor phosphatidylcholine (PC) to the nitrogen of the ethanolamine headgroup of phosphatidylethanolamine (PE) by PE *N*-acyltransferases (NATs) (20). In recent years, more careful characterization of a number of enzymes originally described only as phospholipase A-type enzymes led to their reclassification as NATs. These include phospholipase A2 group 4 epsilon (PLA2G4e) which was shown to be a calcium-dependent NAT and a family of calcium-independent NATs now called phospholipase A/acyltransferase (PLAAT) 1-5 (21). Prior to their reclassification as PLAATs, this family of enzymes had been renamed HRASLS1-5 based on close homology, whereas these enzymes had originally been named A-C1 or Hrev107 (PLAAT1), Adipose-specific phospholipase A (PLAAT3), RARRES3 (PLAAT4), and LRAT-like protein 1 or INAT (PLAAT5). Multiple enzymes can also catalyze the final conversion of NAPEs to NAEs. This conversion can be executed either in a single step by NAPE-hydrolyzing phospholipase D (NAPE-PLD) or in two steps by the serial actions of ABDH4 and GDE1 or an unidentified NAPE phospholipase C and PTPN22 (22, 23).

Vertebrate species whose genomes have been sequenced consistently express multiple NATs as well as the various NAPE hydrolyzing enzymes. For instance, mice express the NATs Plaat1, Plaat3, Plaat5, and Pla2g4e, as well as NAPE hydrolyzing enzymes Napepld, Abdh4, Gde1, and Ptpn22. CRISPR/cas9-mediated gene deletion represents a straightforward method to ablate genes in the NAE biosynthesis pathways, but given the number of genes that potentially must be deleted to identify the gene(s) that control intestinal NAPE biosynthesis, we wanted to utilize an animal model where multiple genes could be deleted relatively quickly and inexpensively. Zebrafish share most of the known signaling mechanism regulating energy balance with mammals, but their small size and high fecundity make them significantly more cost-effective to screen various gene deletions. We therefore performed studies to test if feeding-induced NAE biosynthesis was conserved in zebrafish and the consequence of knocking out various NATs on this feeding response and energy balance.

## MATERIALS AND METHODS

All HPLC grade organic solvents including methanol, chloroform, 1-butanol and acetonitrile were purchased from Fisher (Waltham, MA). Egg yolk powder (cat# E0625), triethylammonium acetate buffer, and methylamine solution (40 wt. % in water) were purchased from Sigma-Aldrich (St. Louis, MO). The internal standards *N*-heptadecanoyl-phosphatidylethanolamine (NHdPE), *N*-[^2^H_4_]heptadecanoyl-ethanolamine ([^2^H_4_]HdEA), and *N*-[^2^H_4_]palmitoyl-PE ( [^2^H_4_]NPPE) were synthesized previously described (24, 25). Other internal standards including *N*-[^2^H_4_]oleoyl-ethanolamine ([^2^H_4_]OEA), and *N*-[^2^H_4_]arachidonoyl-ethanolamine ([^2^H_4_]AEA) were purchased from Cayman Chemicals. Sep-Pak silica cartridges were purchased from the Waters Corporation (Milford, MA). Kinetex 2.6μ C18 100Å column 50×2.1 mm was purchased from Phenomenex (Torrance, CA). Animal studies were approved by the Institutional Animal Care and Use Committee and the Institutional Biosafety Committee of Vanderbilt University.

### Multiple-sequence alignment and phylogenetic tree

Human PLAAT amino acid sequences were used to search for any homologues in zebrafish genome assembly (GRCz11/danRer11) using tblastn. A total of five Plaat isoforms were identified in the zebrafish genome by this method. One had been previously characterized (plaat1), two were previously predicted to be Plaat isoforms (rarres3 and rarres3-like) and two had only been described as unknown proteins (subsequently named plaat1l1 and plaat1l2). Multiple sequence alignment of human, mouse, and zebrafish Plaat isoforms and the phylogenic tree was constructed in EMBL-EBI website using Clustal Omega. The following protein sequences were used to generate this alignment: human PLAAT1, NP_001353041.1; human PLAAT2, NP_060348.1; human PLAAT3, NP_001121675.1; human PLAAT4, NP_004576.2; human PLAAT5, AAH34222.1; mouse PLAAT1, NP_001344021.1; mouse PLAAT3, NP_001349354.1; mouse PLAAT5, AAH61008.1; zebrafish Plaat1, NP_001002590.2; zebrafish Plaat1l1, NP_001116735.1; zebrafish Plaat1l2, NP_001116794.1; zebrafish Plaat4l1, NP_001002623.1; zebrafish Plaat4l2, NP_001070094.2.

### Quantitative real-time PCR

Intestine, liver, brain, and heart of 9 wild-type zebrafish were collected and samples from every three fish were combined to have 3 samples per organs. Samples were homogenized for total RNA extraction using RNeasy Mini Kit (Qiagene). Purified RNAs were reverse transcribed using IScript cDNA Synthesis kit (Bio-Rad). As recommended, samples were treated with RNase-free DNaseI (Qiagene) to remove potential genomic DNA contamination. The RT-qPCRs were performed by using SsoAdvanced Universal SYBR Green Supermix (Bio-Rad). Primers for *plaat1*, *plaat1l1*, *plaat1l2*, *napepld*, and housekeeping gene elongation factor (*elf1a*) used in this experiment are listed in Table 1. Samples were analyzed in duplicate using Bio-Rad CFX96 machine under the following protocol: 95°C for 2 min and 40 cycles of 95°C for 15 sec and 60°C for 30 sec, followed by melt curve analysis of 65-95 °C with 0.5 °C increments at 5 sec/step. Relative gene expression was determined using the ΔCT normalized against housekeeping gene, *elf1a* (*26*).

**Table 1.**
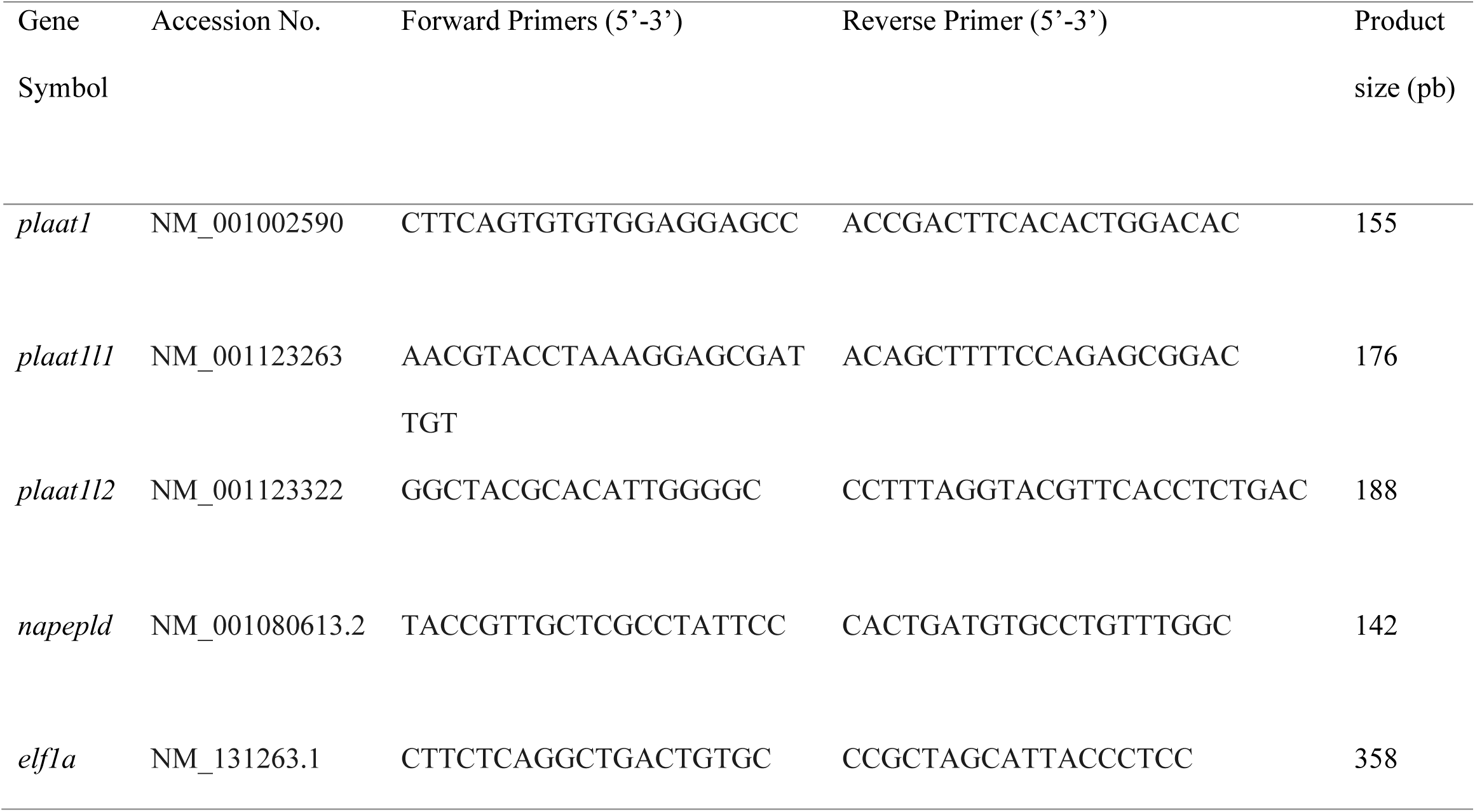
Primer sequences used for quantitative real-time PCR.

**Table 2.**
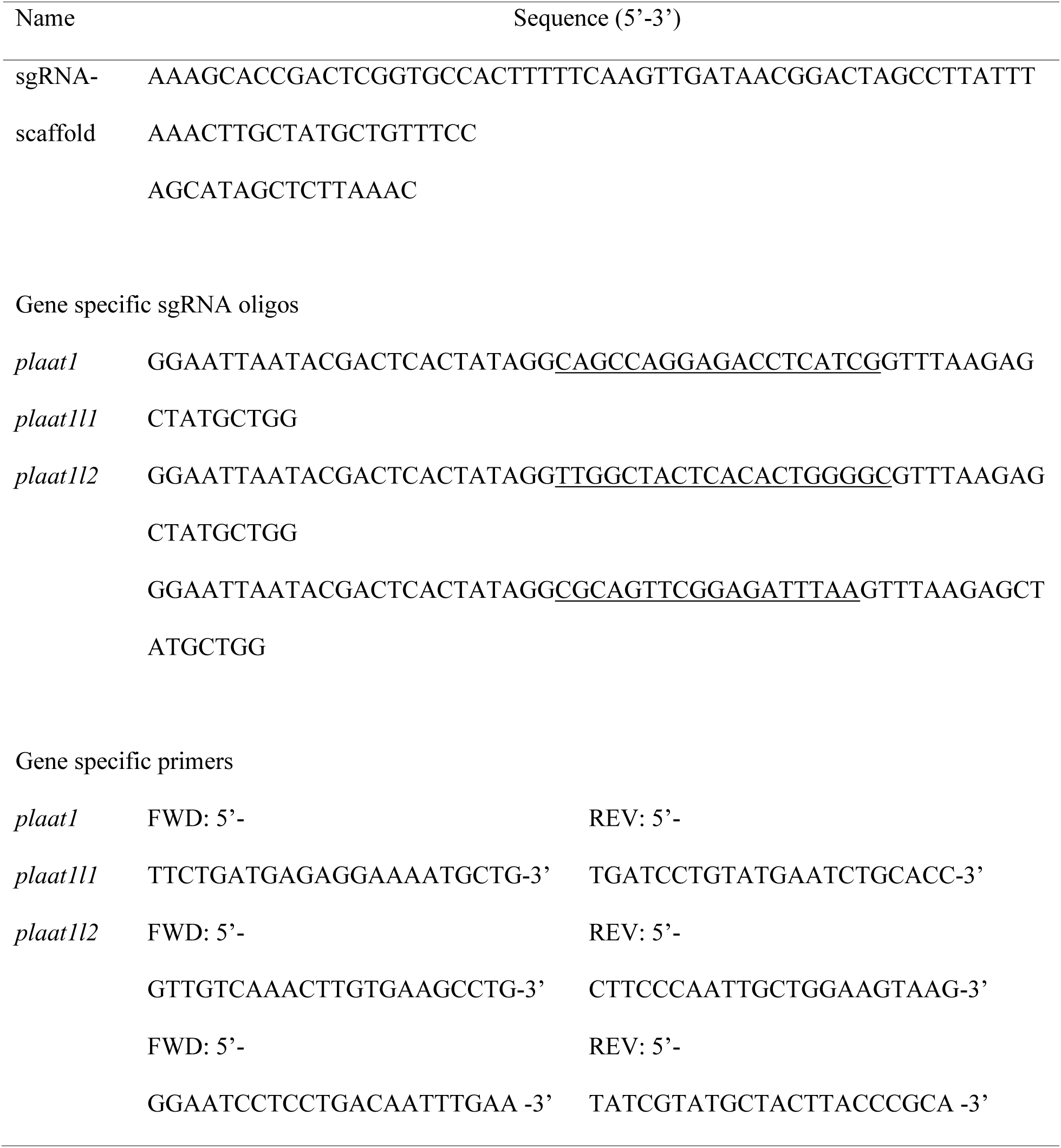
Gene specific sgRNA oligos, primers, and universal oligo containing the sgRNA scaffold (sgRNA-scaffold2).

**Table 3.** Multiple Reaction Monitoring Parameters.

| NAPE | Produced GP-NAE | Precursor ion (m/z) | Product ion (m/z) |
| --- | --- | --- | --- |
| NHdPE | (17:0)GP-NAE | 466.3 | 79.1 |
| [<sup>2</sup>H<sub>4</sub>]NPPE | [<sup>2</sup>H<sub>4</sub>](16:0)GP-NAE | 456.3 | 79.1 |
| *N*-palmitoyl-PE | (16:0)GP-NAE | 452.3 | 79.1 |
| *N*-stearoyl-PE | (18:0)GP-NAE | 480.3 | 79.1 |
| *N*-oleoyl-PE | (18:1)GP-NAE | 478.3 | 79.1 |
| *N*-linoleoyl-PE | (18:2)GP-NAE | 476.3 | 79.1 |
| *N*-eicosapentaenoyl-PE | (20:5)GP-NAE | 498.3 | 79.1 |
| *N*-docosahexaenoyl-PE | (22:6)GP-NAE | 524.3 | 79.1 |

| NAE | Precursor ion (m/z) | Product ion (m/z) |
| --- | --- | --- |
| [<sup>2</sup>H<sub>4</sub>]HdEA | 318.3 | 66.2 |
| [<sup>2</sup>H<sub>4</sub>]OEA | 330.3 | 66.2 |
| [<sup>2</sup>H<sub>4</sub>]AEA | 352.3 | 66.2 |
| PEA | 300.3 | 62.2 |
| SEA | 328.3 | 62.2 |
| OEA | 326.3 | 62.2 |
| LEA | 324.3 | 62.2 |

### Establishing plaat knockout lines

The *plaat1*^-/-^, *plaat1l1*^-/-^, and *plaat1l2*^-/-^ zebrafish lines were generated using sgRNA via CRISPR/cas9 technique as described previously(27, 28). Target-specific sgRNAs were designed using CHOPCHOP (http://chopchop.cbu.uib.no/). The gene-specific oligoes with the identified targets (Table 1. underlined), gene specific primers, and universal oligo containing the sgRNA scaffold primer are listed in Table 1. The double-stranded DNA templates for each sgRNAs were synthesized by mixing 10 pmol of each gene-specific oligo and sgRNA scaffold oligo, 0.5 µl of 10x NEB ThermoPol buffer, dNTPs at final concentration of 200 µM, and 0.5 µl Taq polymerase and using DNase and RNase free water to bring the final volume to 5 µl, followed by PCR under the following cycle: 1 cycle of 94 °C for 1min, 3 cycles of 94 °C 30 sec, decrease to 50 °C at 0.2 °C /sec, 72 °C for 30 s, 1 cycle of 72 °C for 5 min, then cool to room temperature. The MaxiScript T7 kit was used for RNA synthesis as follow: 2 µl 10x T7 transcription buffer, 7 µl RNase free water, 1 µl 10 mM ATP, 1 µl 10 mM CTP, 1 µl 10 mM GTP, 1 µl 10 mM UTP, and 2 µl Enzyme mix was added to the PCR products and incubated at 37 °C for 1 to 2 h. The template DNA was removed by adding 1 µl of TurboDNase I and incubating at 37 °C for another 15 min. The synthesized sgRNAs were purified using mirVANA kit, quantified, and quality checked in a 4% agarose gel.

For mutagenesis, 5 µl of injection mixture consist of 1 µg of recombinant nls-Cas9 (PNA Bio), 200 ng of purified sgRNA, 0.5 µl of 0.5% phenol red was prepared. The mixture was stored at 4 °C overnight for Cas9-sgRNA complex formation. One nL of this mixture was injected directly into the cell of one-cell-stage embryos. The injected embryos were collected in egg water in 10-cm petri dishes. Embryos were raised as founders (F0).

To identify founders with germline mutations, adult F0 zebrafish were crossed to the wild-type fish and their embryos collected at 24 hours post fertilization (hpf). Genomic DNA from eight randomly selected individual embryos from each outcross family was released by alkaline lysis (50 µl of 50 mM NaOH at 95°C for 10 min followed by neutralization with 1M Tris-HCl). Ten µl of lysate from each embryo from the same founder was pools and 1 µl of the mixture was used for PCR using gene specific primers. The heterogeneity of amplicons were evaluated by polyacrylamide gels in heteroduplex mobility assay (HMA) as previously described(28). Pools with mutations were further analyzed using DNA from individual embryos to identify heterozygotes. The PCR product from heterozygotes was Sanger sequenced and the nature of the mutation deciphered manually or by poly peak parser (10.1002/dvdy.24183). Siblings of heterozygotes with a frameshift mutation, preferably with a larger indel, were raised and genotyped at 8 wpf using tailfin DNA and the PCR product sequenced. Heterozygotes with the same desired mutation were raised to adults. They were in-crossed and their offspring (F2) raised and genotyped by PCR using their tailfin DNA. The Sanger sequencing of the PCR product was done by GeneWiz (genewiz.com) . Homozygous F2 were raised to adulthood and used for breeding and performing experiments.

### Effect of plaat genes deletion on whole body NAPEs

Two-month-old wild-type, *plaat1^-/-^, plaat1l1^-/-^, and plaat1l2^-/-^* fish, 3 fish per group, were fasted overnight. To extract their whole-body NAPE, fish were homogenized using Precellys Soft tissue homogenizer CK14 ceramic beads in 1ml 5M NaCl and 0.1 µM final concentrations of internal standards, NHdPE and [^2^H_4_]NPPE. The NAPE extraction and measurement is described below.

### NAPE and NAE measurement in fasted and refed zebrafish

Adult wild-type, *plaat1l1^-/-^, and plaat1l2^-/-^*fish, on a normal diet,14 fish per group, were fasted for 48 h to ensure their gastrointestinal tract is empty. Half of the fish (7 fish) were given Otohime B2, 3.5 mg per fish for 30 min. all the fish were euthanized by tricaine overdose, and their intestines and brain (only for WT fish) were harvested under a light microscope. In the refed fish, most of the food was in the anterior section of their GI, intestinal bulb, and was squeezed out using a pair of tweezers. Samples were placed immediately on dry ice and stored at -80 °C.

The intestinal NAPE measurement experiment was also set up for wild-type zebrafish that are on a high fat diet (HFD) for 6 days. Egg yolk powder was used as the HFD. The fish were transferred to a detached tank with about half litter system water with 30 mg egg yolk powder per fish with occasional stirring. The fish were left in the egg yolk-containing system water for 5 daily for 6 days followed by 48 hours fasting. The rest of the experiment was the same as above.

### NAPE and NAE extraction

Samples (intestines and brains) were sonicated in 700 µl of 5M NaCl and 50 µL of internal standard mixture (10 µL of 10 µM solution of NHdPE, [^2^H_4_]NPPE, [^2^H_4_]OEA, [^2^H_4_]AEA, and [^2^H_4_]HdEA. Ten µL of each sample was taken out for protein measurement. The remainder of each sample was mixed with 300 µL of 5M NaCl, 3 ml of water, and 6 ml of Folch solution (chloroform:methanol 2:1). The mixtures were vortexed for two minutes and left on ice for 1 h with occasional vortexing. Samples were centrifuged at 2000 rpm for 10 min at room temperature. The organic phase, bottom layer, was carefully transferred to a clean 15-ml tube using 1 ml pipet and kept on ice. The aqueous phase was extracted again with 4 ml of chloroform by a brief vortexing. After 30 min on ice, the samples were centrifuged, and the organic phase collected and combined with the first extraction (total of about 8 ml). Samples were dried under N_2_ stream and reconstituted in 1 ml of chloroform. Samples were loaded on prepared Sep-Pak silica columns (Waters) that were previously washed with 4 ml of methanol and equilibrated with 8 ml of chloroform. NAEs were eluted by 8 ml of chloroform:methanol (9:1) followed by elution of NAPEs by Folch solution.

Eluted NAE fractions were dried and reconstituted in 100 µL of chloroform:methanol (1:9). Samples were then filtered using Spin-X 0.22 µm filters (Costar) by centrifuging at 13,000 rpm for 2 min. Filtered samples were transferred to mass-spec vials (Novatech, JGF-30109P-1232).

Eluted NAPE fractions were dried and reconstituted back in 100 µL of methylamination solution (40% methylamine:methanol:1-butanol, 4:4:1). The samples were left in 53 °C water bath for 1 h to convert the NAPE to GP-NAE (glycerophospho-NAE). The samples were then filtered using Spin-X 0.22 µm filters transferred to mass-spec vials.

### NAPEs and NAEs measurement using LC-MS

Samples were analyzed by liquid chromatography (LC) coupled to a ThermoFinnigan Quantum electrospay ionization triple quadrapole mass spectrometer operating in negative ion mode. Samples were injected onto a C18 Phenomenex Kinetex column (2.6 um, 100Å, 50 x 2.1 mm) with the following solvent systems. For NAEs, 0.1% acetic acid in Milli-Q water (mobile phase A) and 0.1% acetic acid in methanol (mobile phase B) were used. Initial conditions were 1% B, with gradient ramp to 99%B over 3 min, held at 99%B for 1 min and then gradient ramp to 1% B in 30 sec, with the column equilibrate at 1% solvent B for 2.5 min prior to next injection. The solvent system for NAPEs was 1 mM triethyl ammonium acetate (mobile phase A) in water and 1 mM triethyl ammonium acetate in acetonitrile (mobile phase B). Initial conditions were 1%B and held for 0.5 min, followed by gradient ramp to 99%B over 3.5 min, held at 99%B for 1.5 min and then gradient ramp to 1%B in 30 sec, with the column equilibrate at 1%B for 2 min prior to next injection.

The mass spectrometer was operated in the multiple reaction monitoring (MRM) mode. The product ion of m/z 62.2 for NAEs and m/z 79.1 for GP-NAEs were used for each specified precursor ion (Table 1 and 2) at a collision energy of 35 eV for NAEs and 50 eV for GP-NAEs. Chromatographic results were analyzed using Xcaliber software (ThermoFinnigan).

### Effect of genotype on body weight trajectory

6 Wild-type and 6 *plaatl1^-/-^* zebrafish, 2.5-month old, were placed on normal feeding of standard diet (TetraMin flakes) 20 mg per fish per day using an automated feeding system (Eheim Automatic Feeding Unit) for 11 weeks. Fish in each genotype were selected so that average body weight per group were similar (130.5±27.6 for WT and 127.3±26.8 for *plaatl1^-/-^*). Body weights were measured every two weeks. One wild-type fish died at week 9 and data from this fish was excluded. At the end of the treatment period, fish were fasted overnight to measure their fasting blood glucose. For this, fish were euthanized following by making an incision just above their caudal peduncle, and collected blood placed directly on blood glucose test strips (Contour, Blood glucose monitoring system).

## RESULTS

### Zebrafish exhibit feeding induced NAPE and NAE biosynthesis

To determine if zebrafish exhibit feeding-induced NA(P)E biosynthesis as reported for rodents, we fasted wild-type (WT) zebrafish for 48 hours and then examined the effect of re-feeding a standard flake diet for 30 minutes on levels of NA(P)Es in the intestinal tract and brain. Individual N-acyl species were measured by LC-MS analysis. In the intestine, total levels of NAPEs (Figure 1A) and NAEs (Figure 1B) increased significantly after re-feeding. Total levels of NAPEs (Figure 1C) and NAEs (Figure 1D) did not significantly increase in the brain after refeeding.

**Figure 1.**
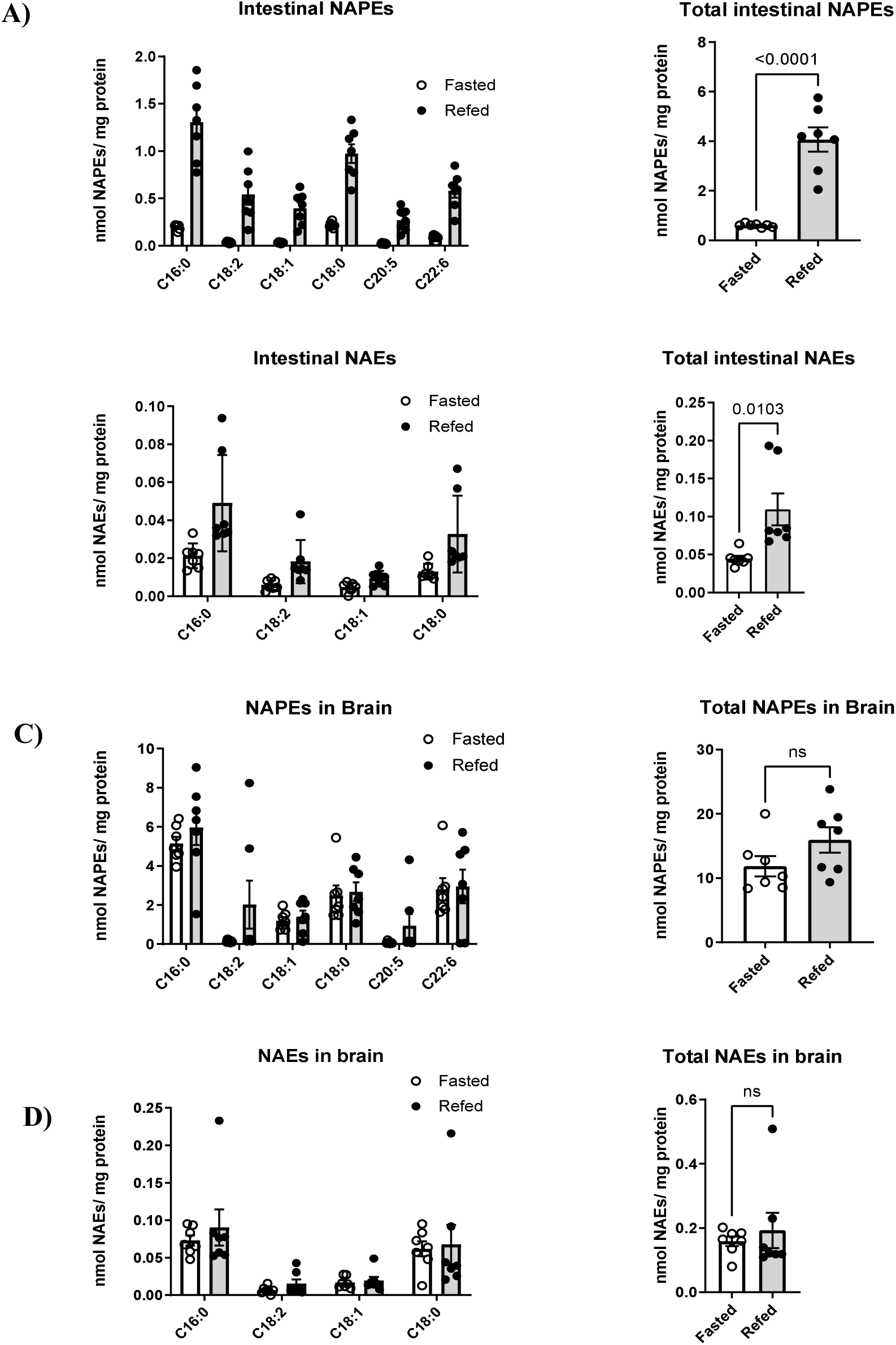
Tissue levels of N-acyl-phosphatidylethanolamine (NAPE) and N-acyl-ethanolamine (NAE) in zebrafish fasted for 48 h and after 30 min of re-feeding with standard flake diet. A) Levels of individual NAPE species (left panel) and total NAPEs (right panel) in whole intestines. B) Levels of individual NAE species (left panel) and total NAEs (right panel) in whole intestine. C) Levels of individual NAPE species (left panel) and total NAPEs (right panel) in whole brain. D) Levels of individual NAE species (left panel) and total NAEs (right panel) in whole brain. All figures are mean ± s.e.m., n=5-7 fish.

### High-fat diet inhibits feeding-induced NA(P)E biosynthesis

To investigate if exposure to high-fat diet (HFD) diminishes feeding-induced NA(P)E biosynthesis in zebrafish similar to what has been shown in rodents, the standard flake diet of WT zebrafish was supplemented with egg yolk powder for 6 days. Fish were then fasted for 48 h and exposed to flake diet or no food for 30 min as before. Fasted fish whose diet had previously been supplemented with the high fat egg yolk showed a significantly diminished rise in their NAPE levels after re-feeding with a standard flake diet compared to fish who had previously received only the standard flake diet (Fig. 2).

**Figure 2.**
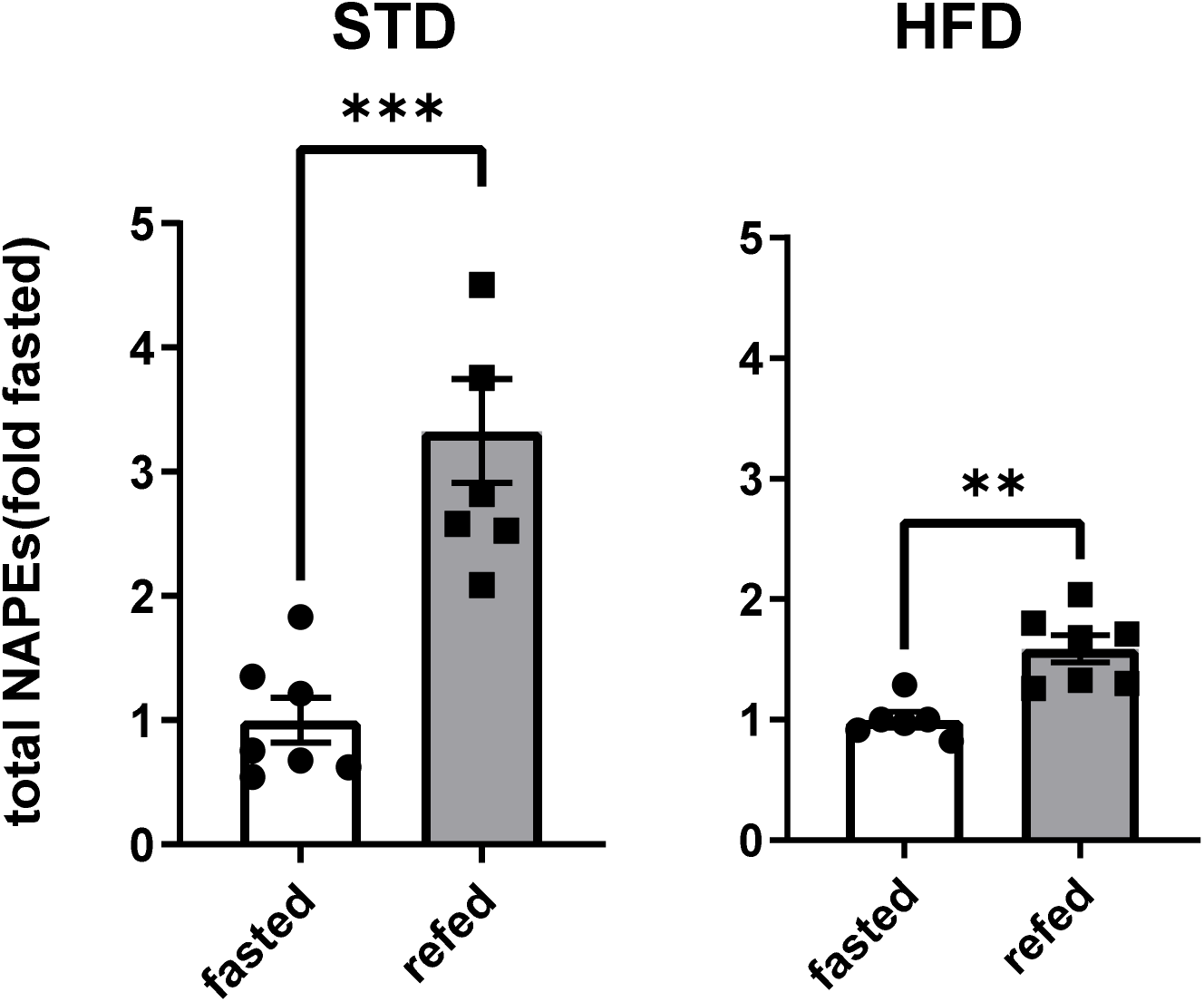
Consumption of high-fat/high cholesterol diet reduces feeding-induced NAPE biosynthesis. Wild-type fish had their standard flake diet supplemented with (HFD) or without (STD) egg yolk for 5 days. Fish were then fasted for 48 h and one half of each group re-fed standard flake diet for 30 min. Total NAPE levels were normalized to the fasted group for each diet. All figures are mean ± s.e.m., n=5-7 fish.

### Identification of *Danio rerio* Phospholipase A/Acyl-transferase (PLAAT) isoforms

We performed a homology search of the zebrafish genome using Clustral Omega using the amino acid sequences of human and mouse PLAAT isoforms as reference. This homology search identified 5 total plaat isoforms, three of which had been previously annotated as such (*plaat1, rarres3/plaat4l1, rarres3l/plaat4l2*) and two that were annotated as unknown proteins (Fig. 3A). These identified genes retained conserved features of the PLAAT catalytic site including appropriately positioned cysteine and histidine residues(^29–35^). The phylogenic tree based on the multiple alignment placed these two new genes most closely to the PLAAT1 family. We therefore named these genes plaat1-like 1 (*plaat1l1*) and 2 (*plaat1l2*) (Fig. 3B). Of note, this version of the phylogenetic tree indicated that *rarres3* is closer to PLAAT2/3 than PLAAT 4/RARRES3.

**Figure 3.**
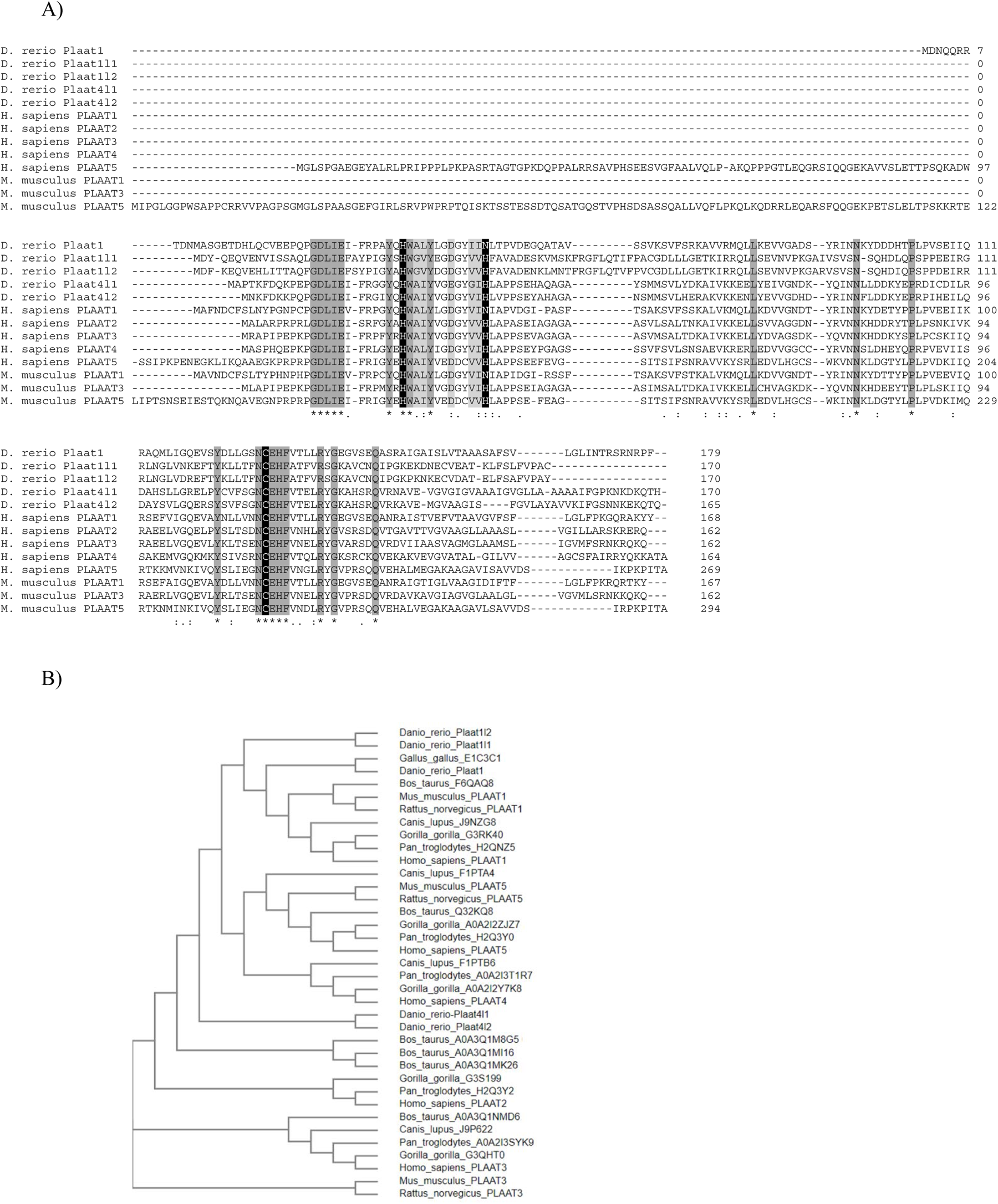
Multiple-sequence alignment and phylogenic tree of zebrafish Plaat isoforms and their homologs in human and mouse. A) Protein alignment generated by EMBL-LBI, asterisks denote residues identical in all PLAAT homologs, with conserved active site residues highlighted with black. B) Calculated phylogenic tree generated by SHOOT (39) using Plaat1l1 sequence for the search input.

### *Plaat1l1* and *plaat1l2* are expressed in the intestinal tract of the zebrafish

We examined the expression of these two novel *plaat* genes, *plaat1l1*, and *plaat1l2*, as well as *plaat1* and the downstream enzyme *napepld*, in key metabolic tissues including intestinal tract, liver, brain, and heart, by quantitative real-time PCR using primers specific to each gene (Table 1). In general, all four genes showed similar patterns of expression, with highest relative expression in the intestinal tract and brain and lower expression in liver and heart (Figure 4). In intestine, the highest *plaat* expression was *plaat1* and *plaat1l1*.

**Figure 4.**
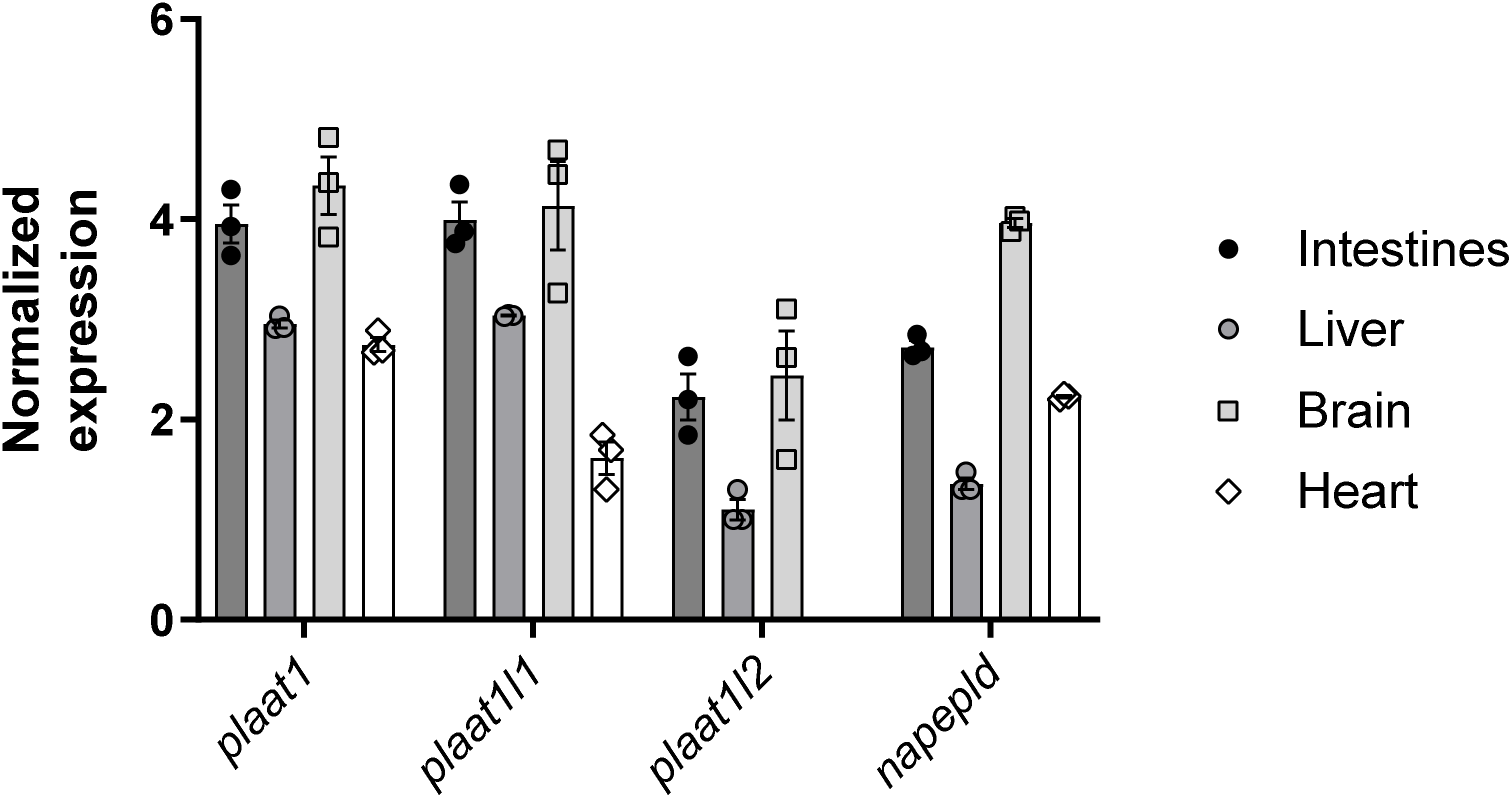
Expression of NAE biosynthetic genes in various zebrafish tissue. qPCR was used to determine the expression of *plaat1*, *plaat1l1*, *plaat1l2*, and *napepld* relative to *elf1a* in wild-type adult zebrafish. N = 3 biological samples, with each biological sample consisting of organs from 3 zebrafish combined.

### Generation of *plaat1*, *plaat1l1*, and *plaat1l2* knock-out zebrafish strains

To determine the contribution of Plaat1, Plaat1l1, and Plaat1l2 to feeding-induced NA(P)E biosynthesis, we generated knock-out strains of each using a CRISPR/Cas approach to generate indels in the early exons of target gene (Fig. 5). The *plaat1^-/-^* strain generated by this process carries a 38 bp frame shift in the first exon that generates a divergent amino acid sequence beginning at residue 28 and that terminates after residue 44, well before the catalytic Cys (Figure 5A). The *plaat1l1^-/-^*strain carries a 22 bp deletion which causes the amino acid sequence to diverge after residue 34 and to terminate after residue 43 (Figure 5B). The *plaat1l2^-/-^*carries a 7 bp deletion which causes the amino acid sequence to diverge after residue 14 and terminate after residue 16 (Figure 5C). To assess whether deletion of these genes reduced levels of NAPEs as anticipated if their products were authentic NAT enzymes, the levels of NAPEs in whole zebrafish of each strain were measured by LC/MS and compared to the levels in wild-type fish. All three knock-out fish showed diminished NAPE levels, with deletion of *plaat1l1* and *plaat1l2* each showing a greater than 50% reduction (Figure 6).

**Figure 5.**
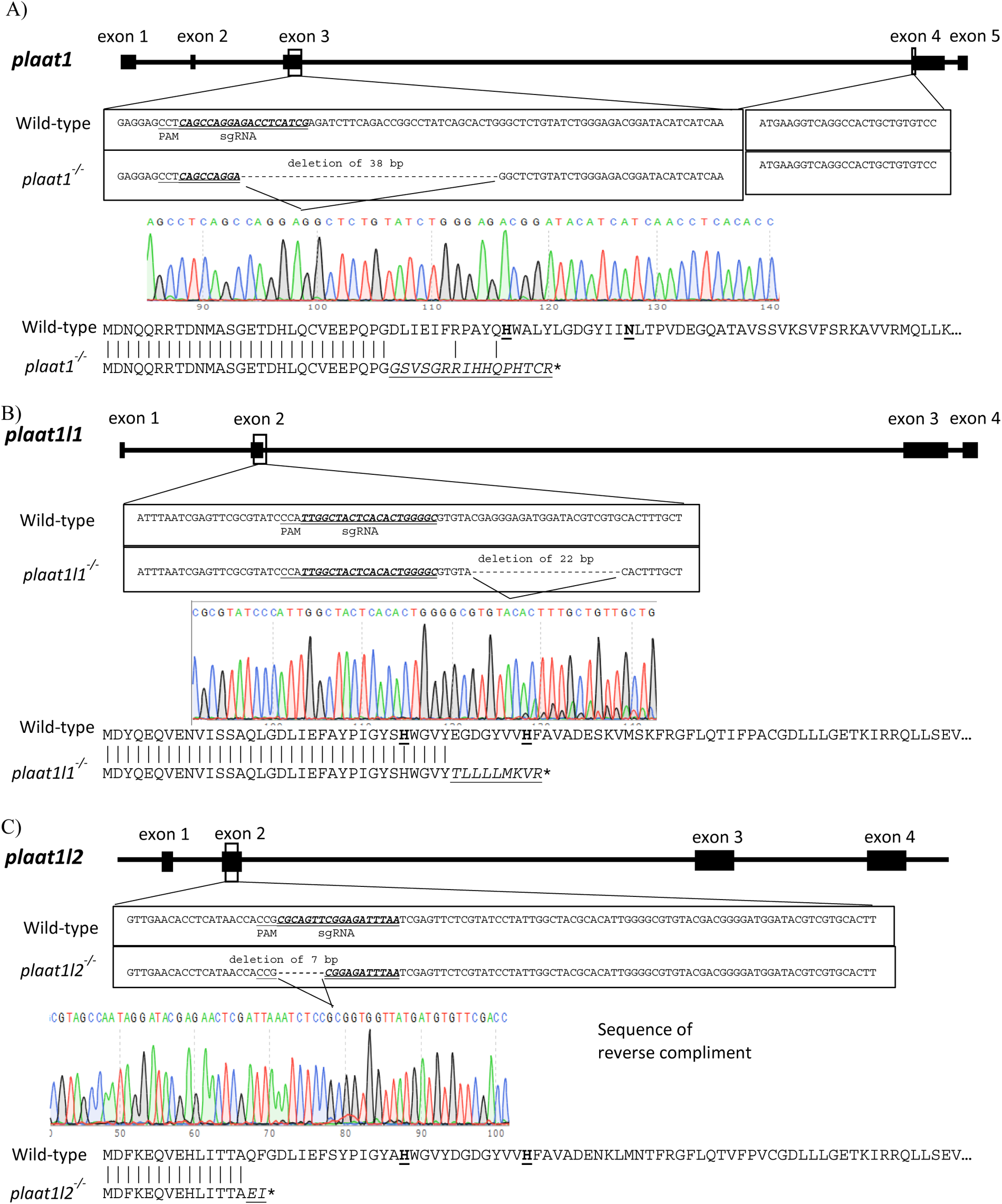
Creation of *plaat* deletion zebrafish lines. CRISPR/cas9 with appropriate sgRNA were used to generate is used to generate the *plaat1*^-/-^, *plaat1l1*^-/-^, and *plaat1l2*^-/-^ zebrafish lines. For each knock-out line, the nucleotide sequence target by the sgRNA is underlined, and the sequencing results, the base pairs deleted in each knockout line, and the predicted change in amino acid, respectively, are shown.

**Figure 6.**
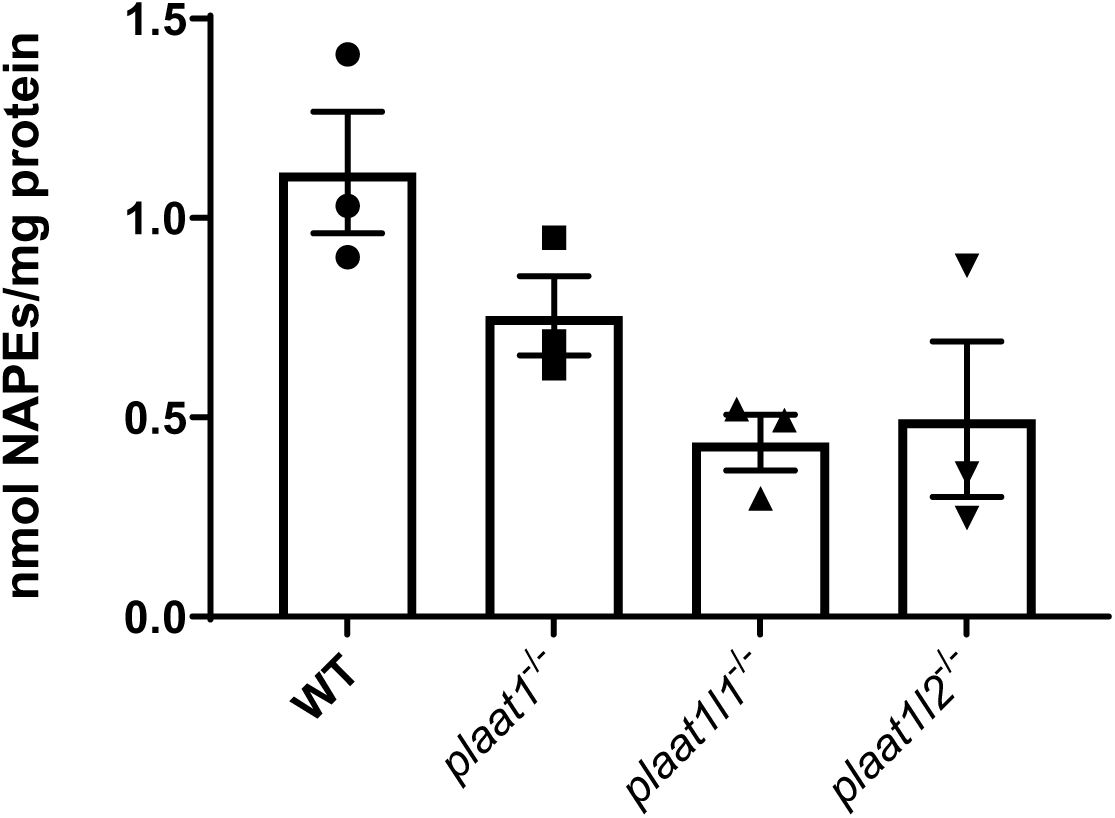
Effect of *plaat* gene deletion on whole body NAPEs. The total NAPEs in whole zebrafish, normalized to total protein, were measured in 2-month-old wild type, *plaat1*^-/-^, *plaat1l1*^-/-^, and *plaat1l2*^-/-^ zebrafish. N = 3 fish per group, results are expressed as mean ± s.e.m.

### Deletion of *plaat1l1* abolishes feeding induced NAPE biosynthesis

Given the significant reductions in whole-body NAPE found for *plaat1l1^-/-^*and *plaat1l2^-/-^*, we examined the effects of these deletions on feeding-induced NA(P)E synthesis. In contrast to wild-type fish that showed significantly increased levels of NAPE after 30 min re-feeding, *plaat1l1^-/-^*fish showed essentially no increase in NAPEs upon re-feeding (Fig. 7). For *plaat1l2^-/-^*fish, fasting levels of NAPEs in *plaat1l2^-/-^* fish were slightly higher than those of either wild-type or *plaat1l1^-/-^* fish and re-feeding significantly increased intestinal NAPE to a level similar to that of re-fed WT fish.

**Figure 7.**
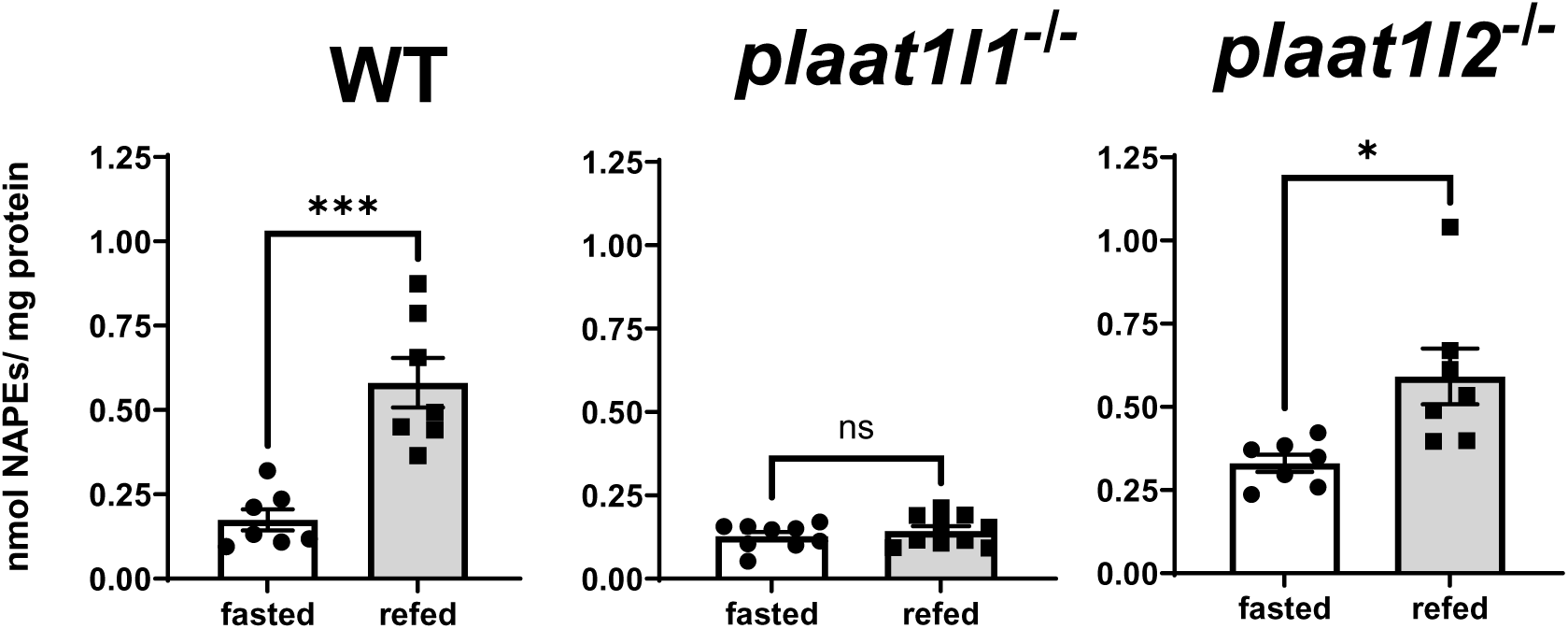
*Plaat1l1^-/-^*fish fail to increase intestinal NAPE levels in response to re-feeding. The levels of NAPE were measured in the intestine of wild-type (left panel), *plaa1tl1*^-/-^ (middle panel), and *plaat1l2*^-/-^ (right panel) zebrafish fasted for 48 hours and half of each group re-fed standard flake diet for 30 min. The results are expressed as total intestinal NAPEs normalized to intestinal protein levels, mean ± SEM, n=7, ***p=0.0003 and *p=0.0120, two-tailed unpaired t-test.

### Deletion of *plaat1l1* causes weight gain, increased BMI, and higher fasted blood glucose in zebrafish on normal diet

To examine the effect of the loss of feeding-induced NA(P)E biosynthesis on whole body energy balance, we compared the weight gain trajectories of a group of wild-type versus *plaat1l1^-/-^* fish that were selected so that they had a similar body weight at day 0 of the study. Body weight was measured at the beginning of the study and then at week 2, 4, 6, 8, and 11. After the final weighing at week 11, fish were euthanized, the length of the fish was determined in order to calculate body mass index and blood collected for glucose levels measurement. Compared to wild-type fish, *plaat1l1^-/-^*fish showed increased weight gain starting by week 2 (Figure 8A). Final body length (measured from snout to tip of caudal peduncle) was similar between the two genotypes (Figure 8B), suggestive that the increased body mass index of *plaat1l1^-/-^* fish (Figure 8C) resulted from greater body fat accumulation rather than faster growth. Blood glucose levels also tended to be higher in *plaat1l1^-/-^* fish versus wild-type fish, consistent with altered glucose handling (Figure 8D).

**Figure 8.**
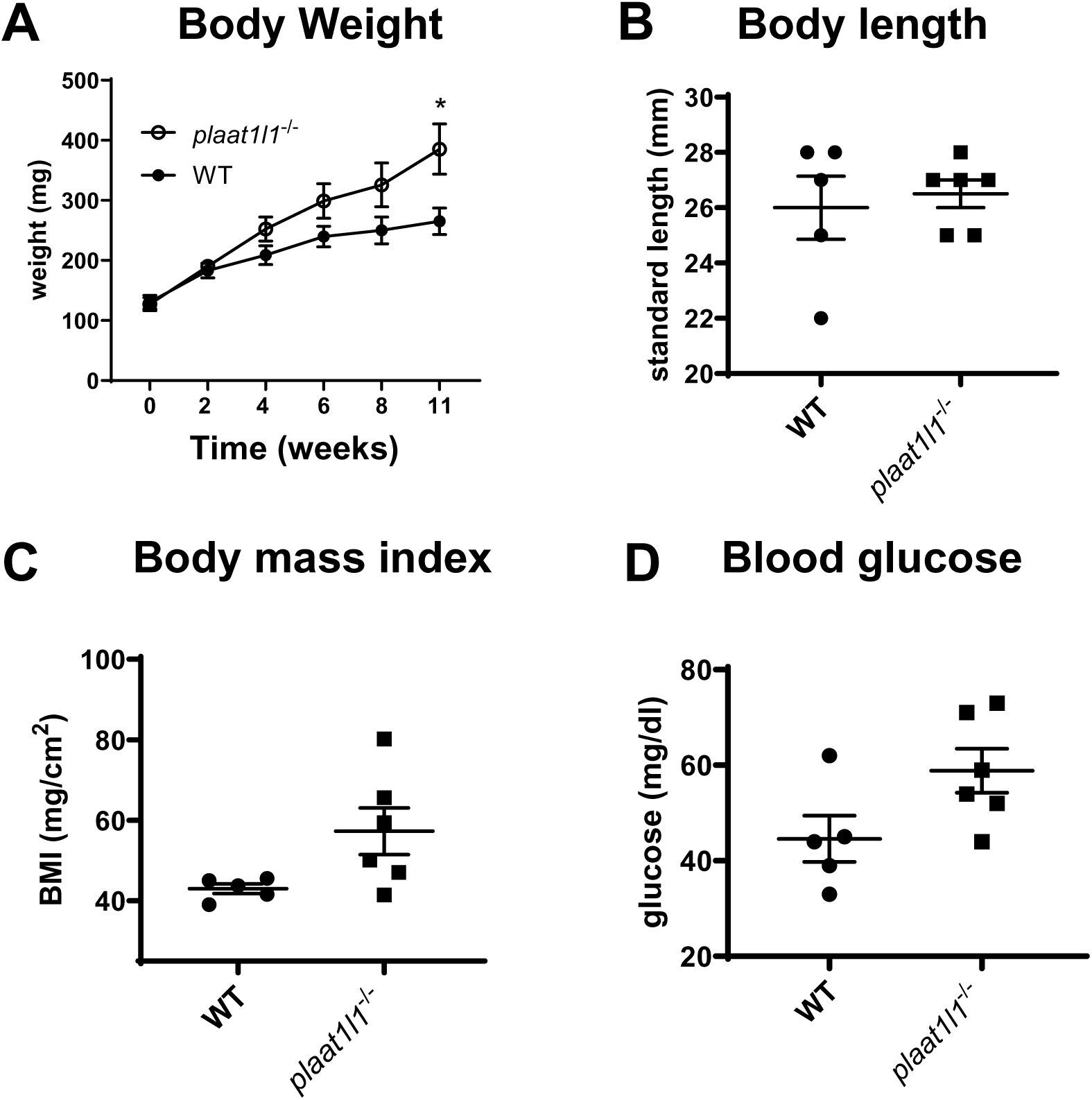
Effect of *plaat1l1^-/-^* on weight gain, body mass index, and fasting blood glucose. Wild-type and *plaat1l1^-/-^* zebrafish of similar weight at day 0 were fed a standard flake diet for 11 weeks and body weight measured periodically. At the end of the 11-week study, fish were euthanized, and body length and blood glucose subsequently measured. A) Body weight, 2-way ANOVA time p=0.0004, genotype p=0.0061, time x genotype p=0.1001, *p=0.0373 uncorrected Fisher’s LSD *wt* vs *plaat1l1^-/-^*week 11. B) Fish length at week 11, p>0.999, two-tailed unpaired t-test. C) Body mass index (mg/cm2) at week 11, p=0.0519, two-tailed unpaired t-test. D) Blood glucose level at week 11, p=0.0632, two-tailed unpaired t-test. All results are presented as mean ± SEM.

## DISCUSSION

Increased intestinal levels of NAEs and their precursors NAPE shortly after consumption of a meal has been demonstrated in rodents and a few other vertebrate species (5–8), but whether this is a physiologically important response for the control of energy balance had not been previously demonstrate and the enzyme(s) responsible for this had not been defined. Here we show that the rapid increase in intestinal levels of NAPE and NAE in response to feeding seen in rodents is conserved in zebrafish, that high fat diet also significantly reduces this response in zebrafish, that the calcium-independent NAT plaat1l1 is responsible for generating the increase in intestinal NAPEs (and thus NAEs), and that fish lacking the ability to acutely increase intestinal NAPE levels in response to feeding have increased weight gain.

The rapid rise in intestinal NAEs and their precursors NAPE after feeding had previously been postulated to regulate energy balance, based on the finding that pharmacological administration of OEA and other NAEs (except AEA) reduce food intake, increase fatty acid oxidation, and reduce weight and fat gain in rodents (11, 17), as well as the finding that this response was lost in rodents fed an obesogenic diet (10, 17). However, that feeding-induced biosynthesis of NAEs was crucial to appropriate energy balance had not been definitively demonstrated. Attempts to address the physiologic importance of this response by intestinal specific ablation of *Napepld*, an enzyme that directly hydrolyzes NAPEs to NAEs, in mice were inconclusive. Mice with intestinal specific deletion of Napepld (Napepld^ΔIEC^ mice) show slightly lower fasting levels of OEA (but not other NAEs) in the jejunum (36). When fasted mice that were previously only exposed to normal diet were given their first high-fat meal, this tended to raise OEA levels in both wild-type and Napepld^ΔIEC^ mice 60 min after initial intake of the meal (36). Subsequent studies suggested that there might be a delay in the rise in jejunal OEA in Napepld^ΔIEC^ mice, as wild-type but not Napepld^ΔIEC^ mice showed a trend to increased jejunal OEA levels 30 min after exposure to fat, while both showed trends to increases at 60 min (37). Given the limited effects of Napepld deletion on jejunal OEA levels after fat feeding, interpreting the effects on energy balance is challenging. The body weight of Napepld^ΔIEC^ mice did not differ from wild-type mice when both were fed normal chow diet (36); however, Napepld^ΔIEC^ mice chronically fed a high-fat diet did show increased body weight and adipose weight compared to wild-type mice. Thus, even minor reductions in the intestinal biosynthesis of NAEs in response to feeding might impact energy balance long-term when additional stressors are applied. Napepld^ΔIEC^ mice did not show any significant changes in glucose handling compared to wild-type mice (36).

Because the biosynthesis of NAPEs was previously shown to be the rate limiting step in NAE production (19), we reasoned that genetic ablation of the enzymes producing NAPEs could potentially produce a more profound reduction in intestinal NAE biosynthesis in response to food intake than *Napepld* deletion. Given that multiple PLAAT isoforms and PLA2G4E could be responsible for feeding induced NAPE synthesis, we turned to an animal model where rapid deletion of multiple genes would be feasible. Because of their high fecundity, ease of genetic manipulation, and similarity to mammalian metabolism, zebrafish (*Danio rerio*) has been used for genetic analysis of many physiologic processes including energy homeostasis (38). Our analysis of the zebrafish genome for sequence homology with mammalian PLAATs identified two genes as likely *plaat* isoforms that had been previously annotated simply as genes of unknown function. Because phylogenic analysis suggested these two genes were most closely related to *plaat1/PLAAT1/Plaat1*, we named them *plaat1l1* and *plaat1l2*. Importantly, Plaat1l1 and Plaat1l2 share the appropriately placed catalytic triad residues (His, Cys, Asn/His) and the conserved NCEHF(V/A) found in all other PLAATs. We then evaluated the contribution of these enzymes to feeding-induced NAPE biosynthesis. We found that deletion of *plaat1l1* completely ablated the increase in intestinal NAPE and NAEs seen by 60 min after food intake in wild-type fish. These results suggest that a PLAAT isoform and not PLA2G4E is likely to be the rate-limiting enzyme in feeding-induced NAE biosynthesis in most species. The identification of *plaat1l1* as the NAT responsible for feeding-induced NAPE biosynthesis should enable future studies to determine the exact mechanisms for this response and why high-fat diets ablates this response.

Our finding that fish that lack the ability to biosynthesize NAPEs (and therefore NAEs) in response to feeding also have increased weight gain and poorer glucose handling, supports that notion that feeding-induced NAPE and NAE biosynthesis is critical for appropriate regulation of energy balance, so that the loss of this biosynthesis that occurs with consuming a high fat diet in both rodents and zebrafish is likely to be a contributing factor as to why chronic consumption of high fat diets result in obesity, insulin resistance, and type 2 diabetes. This finding suggests that therapeutic interventions either to preserve or rescue PLAAT-dependent NAPE biosynthesis in the face of obesogenic diets may be a valuable approach to cardiometabolic diseases.

## Funding

This study was supported by faculty funds from the Vanderbilt University Department of Pharmacology.

## Abbreviations

NAFLD: non-alcoholic fatty liver diseases
NAPE: N-acyl-phosphatidylethanolamine
NAT: N-acyltransferase
NAE: N-acyl-ethanolamine
PEA: N-palmitoyl-ethanolamine
SEA: N-stearoyl-ethanolamine
OEA: N-oleoyl-ethanolamine
LEA: N-linoleoyl-ethanolamine
HFD: high fat diet
PC: phosphatidylcholine
PE: phosphatidylethanolamine
PLAAT: phospholipase A/acyltransferase
Hpf: hours post fertilization

## Data Availability

Processed data generated during the current study are included in the published article. All raw and processed data for these studies are archived at the FigShare repository Web site 10.6084/m9.figshare.24871527.

## Acknowledgement

The authors would like to thank the staff of the Vanderbilt Zebrafish Core and the Vanderbilt Mass Spectrometry Core for technical assistance. We also thank Ms. Isabelle Suero and Mr. Andrew Jenkins for assistance with zebrafish breeding and maintenance.

## Notes

https://figshare.com/10.6084/m9.figshare.24871527

### Competing Interest Statement

The authors have declared no competing interest.

